# Methylated PP2A stabilizes Gcn4 to enable a methionine-induced anabolic program

**DOI:** 10.1101/2020.05.05.079020

**Authors:** Adhish S. Walvekar, Ganesh Kadamur, Sreesa Sreedharan, Ritu Gupta, Rajalakshmi Srinivasan, Sunil Laxman

## Abstract

Methionine, through S-adenosylmethionine, activates multifaceted growth programs where ribosome biogenesis, carbon metabolism, amino acid and nucleotide biosynthesis are induced. This growth program requires activity of the Gcn4 transcription factor (called ATF4 in mammals), which enables metabolic precursor supply essential for anabolism. Here, we discover how the Gcn4 protein is induced by methionine, despite conditions of high translation and anabolism. This induction mechanism is independent of transcription, as well as the conventional Gcn2/eIF2α mediated increased translation of Gcn4. Instead, when methionine is abundant, Gcn4 ubiqitination and therefore degradation is reduced, due to the decreased phosphorylation of this protein. This Gcn4 stabilization is mediated by the activity of the conserved methyltransferase, Ppm1, which specifically methylates the catalytic subunit of protein phosphatase PP2A when methionine is abundant. This methylation of PP2A shifts the balance of Gcn4 to a dephosphorylated state, which stabilizes the protein. The loss of Ppm1, or PP2A-methylation destabilizes Gcn4 when methionine is abundant, and the Gcn4-dependent anabolic program collapses. These findings reveal a novel signaling and regulatory axis, where methionine directs a conserved methyltransferase Ppm1, via its target phosphatase PP2A, to selectively stabilize Gcn4. Thereby, when methionine is abundant, cells conditionally modify a major phosphatase in order to stabilize a metabolic master-regulator and drive anabolism.

## Introduction

Cellular commitments to different states, such as growth, survival, or self-destruction depend on multiple cues. The availability of nutrients is a critical signal for cells to switch to a proliferative state. When adequate supplies of carbon and nitrogen are available, cells channel them into energy production, biomass generation and ribosomal biogenesis, all leading to cell growth and division. Indeed, using simple eukaryotic model organisms like yeast, several comprehensive, systems-level studies have investigated responses of cells to distinct nutrient availabilities, to define transcriptional programs that indicate growth or starvation states (1–6). When cells commit to mitotic division, multiple signalling cascades and transcriptional responses control overall ‘growth programs’ (1, 3, 6–10). While global responses to nutrient changes are well documented, specific signalling and regulatory mechanisms that directly couple the sensing of specific metabolites to metabolic programs remain more poorly studied. Thus, there is a need to identify these regulatory mechanisms, in order to mechanistically understand growth programs.

In this regard, some metabolites directly act as “sentinel” molecules that trigger growth programs. One such metabolite that signals growth programs is methionine (11). Studies using yeast cells revealed that methionine strongly inhibits autophagy, and concomitantly also increases cell proliferation (12–14). Methionine (via its metabolism to S-adenoyslmethionine, SAM) also activates the Target of Rapamycin (TOR) signaling in yeast and mammalian cells (13, 15, 16), regulates lipid balance, increases translational capacity and maintains metabolic homeostasis during growth (17). Consistent with these roles, studies from a variety of cancers also suggest that methionine is a key determinant of tumor cell proliferation (18–21). Indeed, a fundamental determinant of cell growth is the steady supply of anabolic precursor molecules such as amino acids and nucleotides, which fuel sufficient translation and RNA/DNA synthesis. Methionine is itself incapable of being a metabolic precursor to these building blocks, and instead appears to function as a signaling molecule, activating these processes, likely via its primary metabolite s-adenosyl methionine. However, specific signaling events that directly control this metabolic transformation leading to the continuous supply of anabolic molecules remain unclear.

In a recent study, we discovered that methionine controls a hierarchically organized, well defined anabolic program (14). The addition of methionine induces the expression of genes involved in translation and ribosomal biogenesis, as well as key metabolic reactions, primarily the pentose-phosphate pathway, glutamine synthesis, and amino acid and nucleotide biosynthesis. In short, addition of methionine switched cells to an anabolic state and enabled efficient assimilation and utilization of available carbon and nitrogen sources, resulting in increased amino acid and nucleotide biosynthesis required for growth (14, 22). Surprisingly, in order to sustain the growth program, cells required the activity of what is primarily considered a starvation/survival response factor, Gcn4, to balance metabolism with translation (22).

Gcn4 (called ATF4 in mammals), is a well-studied transcriptional activator of genes in amino acid biosynthetic pathways (23, 24). Gcn4 function has primarily been studied under amino acid starvation conditions, where it has a key role in restoring amino acid homeostasis (24–27). When cells are starved even of one amino acid, the translation of Gcn4/ATF4 transcripts is enhanced in a Gcn2/phospho-eIF2α dependent manner. During amino acid starvation, the Gcn2 protein kinase is activated, downregulating global translation with concomitant enhancement of Gcn4 translation. This promotes amino acid biosynthetic gene expression, in order for cells to restore amino acid pools (24). Consequently, Gcn4/ATF4 is annotated as a marker of the cellular amino acid starvation response. Paradoxically, our earlier findings (14, 22) are noteworthy because Gcn4 was critical in a context of high growth and proliferation, rather than survival, when methionine alone was supplemented. Therefore, in such a context of *increased* proliferation, what signaling processes control Gcn4 induction? This remains unknown.

In this study, we elucidate a novel mechanism that increases Gcn4 protein during this growth program. Specifically, abundant methionine does not additionally enhance Gcn4 transcription or translation. Instead, methionine post-translationally results in increased Gcn4 protein, via a SAM-dependent methylation of the protein phosphatase PP2A by the methyl transferase Ppm1 (LCMT1 in mammals). Methylated-PP2A maintains Gcn4 in a dephosphorylated state, preventing its subsequent degradation which typically occurs through a phosphorylation dependent ubiquitination of Gcn4. The stabilization of Gcn4 is not observed in the absence of Ppm1, or the loss of PP2A methylation. Further, PP2A methylation mediated stabilization of Gcn4 is required for high *de novo* amino acid and nucleotide biosynthesis in abundant methionine. Our findings illustrate how a methionine/SAM responsive protein phosphatase (PP2A) directly controls anabolism, by increasing amounts of a metabolic master-regulator, Gcn4. This reveals how a methionine-responsive signaling response can coordinately control the metabolic state of the cell.

## Results

### Gcn4 is induced by methionine while cells retain a high translation state

In yeast cells growing in overall amino acid limited conditions, supplementing methionine drives a transcriptional and metabolic growth program (14, 22). Methionine transcriptionally and functionally induces rate-limiting reactions in amino acid biosynthesis, leading to enhanced amino acid and nucleotide pools (14). Methionine also transcriptionally induces ribosomal genes, suggesting that it upregulates overall protein synthesis. To test if this transcriptional induction of ribosomal genes was functionally observed as increased translation, we examined the impact of methionine addition on global translation using polysome profiling, with cells in rich medium (yeast extract, peptone, 2% lactate) as a control. Expectedly, yeast grown in a complex, rich medium (**RM**) showed a 3:1 polysome:monosome ratio indicating robust translation in the system. Contrastingly, cells shifted to a defined minimal medium with the same carbon source (**MM**) showed a drastic reduction in the polysome pool and an increased monosome peak (polysome:monosome ratio of ∼1:1), reflective of low translation. Methionine supplementation during a shift to MM medium (**MM+Met**) significantly restored polysome levels, resulting in a polysome:monosome ratio of ∼1.7:1 (Figure 1A). This increase in polysome levels is greater than that observed with supplementing all non-sulphur amino acids combined except tyrosine (pool of 17 amino acids, excluding methionine and cysteine). These data therefore confirm that methionine supplementation increases global translation in otherwise amino acid limited conditions, consistent with the expectations from the observed anabolic program.

**Figure 1:**
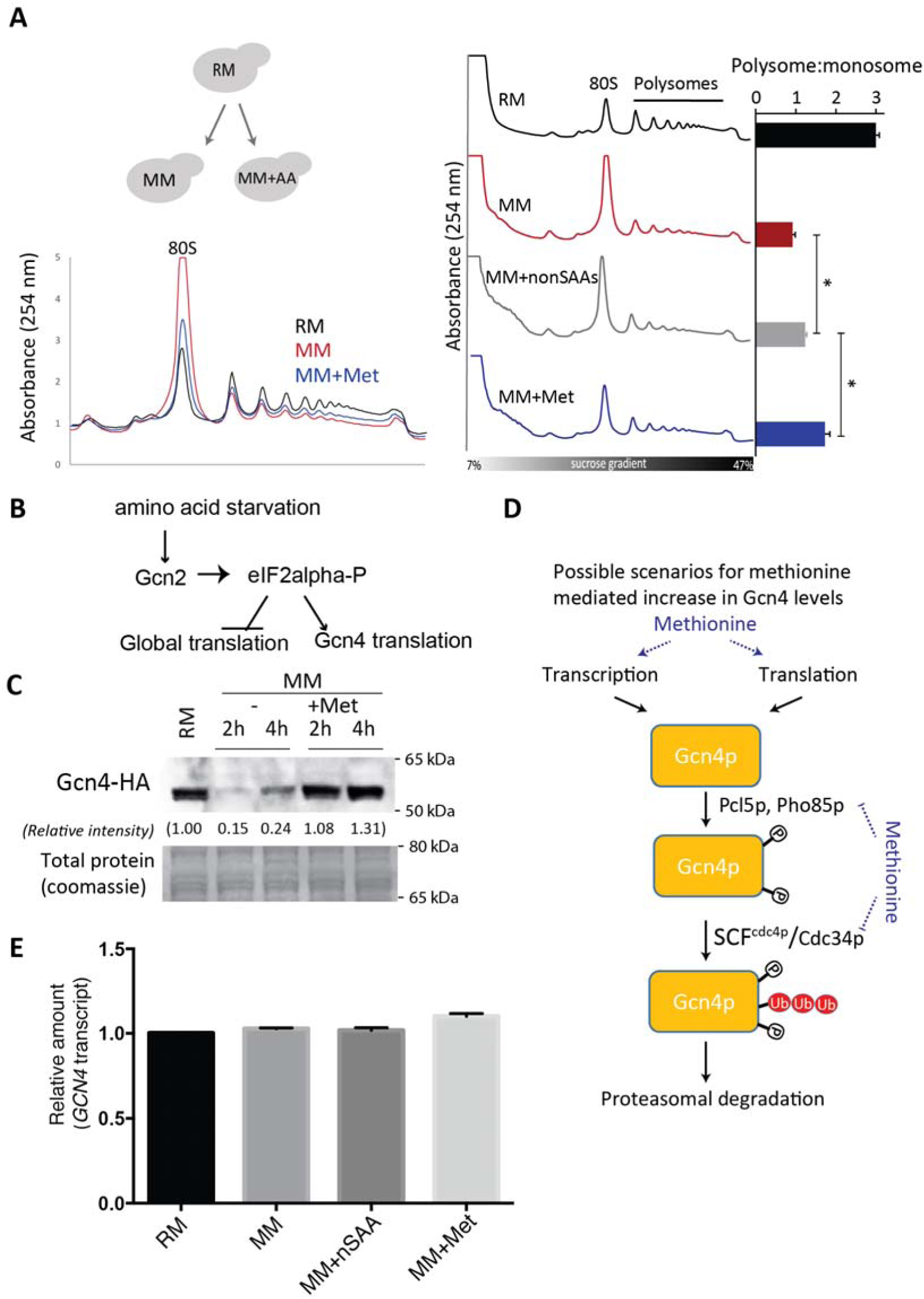
Gcn4 is induced by methionine while cells retain a high translation state. A) Polysome profiles and polysome:monosome ratios of cells growing in rich medium (RM), minimal medium (MM), or minimal medium supplemented with methionine (MM+Met). Plots shown are from three experiments, mean ± SD. * p<0.05 (Student’s t-test). Also see Supplemental Figure 1. B) A canonical model of Gcn4 regulation, mediated via increased translation of Gcn4 transcripts, controlled by the activation of the Gcn2 kinase and the phosphorylation of eIF2α. C) Methionine-dependent Gcn4 induction: Gcn4 protein amounts in cells growing in RM, MM or MM+Met, as measured by a western blot. The blot shown is representative of independent experiments done at least 4 times. D) Relative amounts of Gcn4 transcripts in RM, MM and MM+Met. Data shown are from whole-transcriptome (RNA-seq) data from biological replicates, obtained from (14). Differences in Gcn4 transcript amounts between any two of the different conditions are statistically not significant (Student’s t-test).

While dissecting the anabolic program that methionine controls, we had unexpectedly uncovered a critical role for Gcn4 (14, 22). Gcn4 is a transcription factor that has been best studied as an activator of amino acid biosynthetic genes during severe amino acid starvation, to restore intracellular amino acid homeostasis (24–26, 28–30). During starvation conditions (including methionine starvation), the Gcn2 kinase phosphorylates eIF2α, resulting in a damping of global translation with simultaneous increase in Gcn4 levels (Figure 1B) (24, 31). Our observations in methionine replete conditions therefore posted a paradox. Here, global translation is clearly increased (Figure 1A), yet Gcn4 function appears to be critical under these conditions. Hence, we systematically addressed how abundant methionine might regulate Gcn4.

Using a strain in which a C-terminal hemagglutinin (HA) tag was inserted at the chromosomal GCN4 locus, we measured steady state levels of Gcn4 protein after a shift from RM to MM. As observed earlier (14, 22), Gcn4 levels were high only in cells shifted from robust growth conditions (RM) to methionine supplemented medium (MM+Met), in contrast to cells shifted to medium with no amino acids (MM) (Figure 1C), or even with all non sulphur amino acids supplemented except tyrosine (MM+nonSAA), as shown earlier (14). This reiterated the above stated paradox; when methionine is abundant, global translation increases, but Gcn4 amounts are strongly induced as well. Therefore, how might abundant methionine increase Gcn4 levels?

### Gcn4 accumulation due to methionine does not depend on the GCN2-translation axis

Gcn4 protein levels can be fine-tuned through possible transcriptional, translational or post-translational mechanisms (Figure 1D). We investigated which of these steps is under methionine regulation. For this, we first compared mRNA levels of GCN4 in medium with and without methionine supplementation, using RNAseq datasets from our previous study (14). GCN4 transcript levels were only marginally higher in MM with methionine than in medium with all amino acids except methionine/cysteine, or minimal medium without any amino acid supplementation, and this difference was not significant (Figure 1E). Thus, GCN4 expression is not significantly transcriptionally regulated by methionine.

To test whether Gcn4 is actively synthesized after methionine addition, we treated cells with cycloheximide, a commonly used ribosome blocker. Gcn4 amounts were reduced when cells were shifted to methionine replete medium supplemented with cycloheximide (Figure 2A). This expectedly indicates that Gcn4 is continuously, actively translated under these conditions.

**Figure 2:**
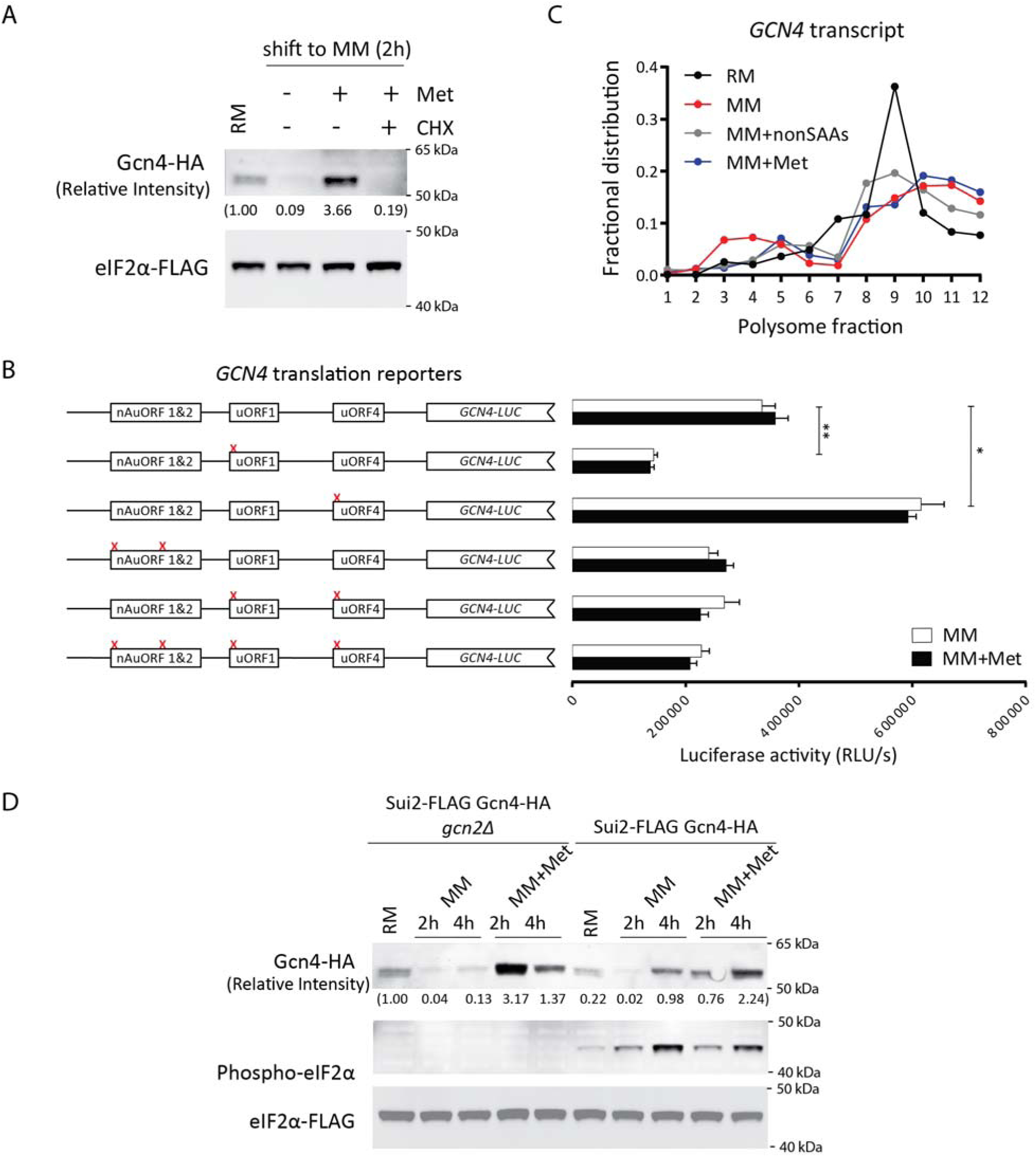
Methionine induced Gcn4 accumulation does not depend on the GCN2-translation axis. A) Effect of cycloheximide treatment on Gcn4 protein amounts. Cells growing in MM or MM+Met were treated with cycloheximide and Gcn4 protein amounts were estimated by Western blotting. The gel is representative of at least three biological replicates. B) Relative *GCN4* translation in MM or MM+Met, as measured using a series of luciferase-based *GCN4* translation reporters. The relative luciferase activity is shown on the y-axis of the plots, while the different reporters used are illustrated on the left. The data shown are from three biological replicates, mean ± SD. * p<0.05 (Student’s t-test). Also see Supplemental Figure 1. C) *GCN4* transcript amounts in different polysome fractions, obtained from cells grown in RM, MM or MM+met. Transcripts were measured using standard, quantitative RT-qPCR approaches. Also see Supplemental Figure 2A. D) Gcn4 protein amounts estimated in wild-type or *gcn2Δ* cells, grown in MM or MM+Met. The yeast ortholog of eIF2α is Sui2, which was tagged with a FLAG-epitope (in the native chromosomal locus). As controls, the amounts of phosphorylated eIF2α and total eIF2α protein are also shown.

The post-transcriptional regulation of Gcn4 is complex and achieved by multiple modes (24). The most extensively studied regulation is through the control of *GCN4* translation during amino acid starvation. During starvation the enhanced translation of *GCN4* transcripts is mediated by the activated Gcn2 kinase via phosphorylated eIF2α (24, 32, 33). Here, *GCN4* translation is dictated by its 5′ leader sequence (Figure 2B), primarily by the upstream open reading frame 1 (uORF1) and uORF4, as well as through two non-AUG uORFs (nAuORFs) that are present upstream of the uORFs (24, 34–40). We investigated these mechanisms of increased translation of *GCN4*, in a methionine-dependent context, by comparing *GCN4* mRNA translation in cells growing in MM medium with or without methionine supplementation. We used a standard *GCN4*-luciferase fusion reporter that indicates *GCN4* translation (40, 41) to ask if these features also contribute to potential methionine-specific enhancement of *GCN4* translation. A series of *GCN4*-luciferase translation reporters with point mutations in each regulatory element were included, and were all validated using a conventional method of *GCN4* translation activation by adding rapamycin (Supplementary Figure 1A and 1B). Using these reporters, we compared translation of *GCN4* in minimal medium with or without methionine supplementation (Figure 2B), or in medium with all amino acids except methionine and cysteine (Supplementary Figure 1C). Notably, this reporter activity was similarly high in cells shifted to MM, both with and without methionine supplementation (Figure 2B). This indicates continuing, high translation of the GCN4 transcript in both these conditions (MM and MM+Met). Importantly, the luciferase reporter activity was not significantly further enhanced in the presence of methionine, indicating that the translation rate of the *GCN4* ORF is persistently high in minimal medium regardless of methionine addition. To independently validate this, we measured the distribution of native *GCN4* transcript in polysome fractions of cells shifted to MM medium with or without amino acid supplementations. Again, we found no notable change in the *GCN4* transcript distribution in any of the combinations tested (Figure 1C and Supplementary Figure 1D). Collectively, our data suggest that GCN4 is actively translated at comparable rates in MM medium regardless of methionine supplementation.

These data suggested that the resulting Gcn4 accumulation in methionine might be independently regulated beyond the GCN2-eIF2α regulatory axis. To test this, we asked whether the increased Gcn4 amounts seen in the presence of methionine depended on GCN2. Even in a *gcn2*Δ strain, Gcn4 accumulated upon methionine supplementation (Figure 2D), suggesting that other mechanisms contribute to the methionine-mediated accumulation of Gcn4. Finally, we compared cell proliferation in MM+Met, in wildtype, *gcn2*Δ and *gcn4*Δ cells. Notably, in this condition, *gcn2*Δ grew significantly better (partial rescue) than *gcn4*Δ cells in MM+Met (Supplementary Figure 2). Here, this decrease in growth, as well as the decrease of Gcn4 in the *gcn2*Δ strain is consistent with the role of Gcn2 in maintaining synthesis, but not Gcn4 stability or turnover. Therefore, the canonical mechanism of *GCN4* control at the translational level by the GCN2-eIF2α axis cannot explain the increased Gcn4 levels specifically in the presence of methionine, suggesting the presence of alternate post-translational mechanisms in regulation of Gcn4 levels.

### Methionine inhibits Gcn4 degradation by the 26S proteasome

Regulation of Gcn4 has been extensively investigated at the level of its synthesis, but also occurs at the level of degradation (Figure 1D). Targeted degradation of Gcn4 involves its polyubiquitination and subsequent degradation by the 26S proteasome. We therefore asked whether methionine modulates proteasomal degradation of Gcn4. We devised a modified experimental setup where the role of methionine in Gcn4 degradation could be specifically examined (Figure 3A). We first allowed Gcn4 to accumulate in MM+Met, and then measured protein levels after shifting the cells to medium lacking methionine. Indeed, Gcn4 protein levels rapidly decreased even within 20min of shift of cells to MM (Figure 3A), or a shift to MM+nonSAAs (Figure 3B), consistent with a possible role for methionine in stabilizing Gcn4 by modulating its degradation. We next tested whether the decrease in Gcn4 amounts observed after methionine was removed could be reversed in the presence of MG-132, a commonly used proteasome inhibitor. Here, the rapid degradation of Gcn4 was inhibited by addition of MG-132 to MM or MM+nonSAAs (Figure 3C and Figure 3D). These results suggest that in amino-acid limited conditions, the availability of methionine inhibits Gcn4 degradation.

**Figure 3:**
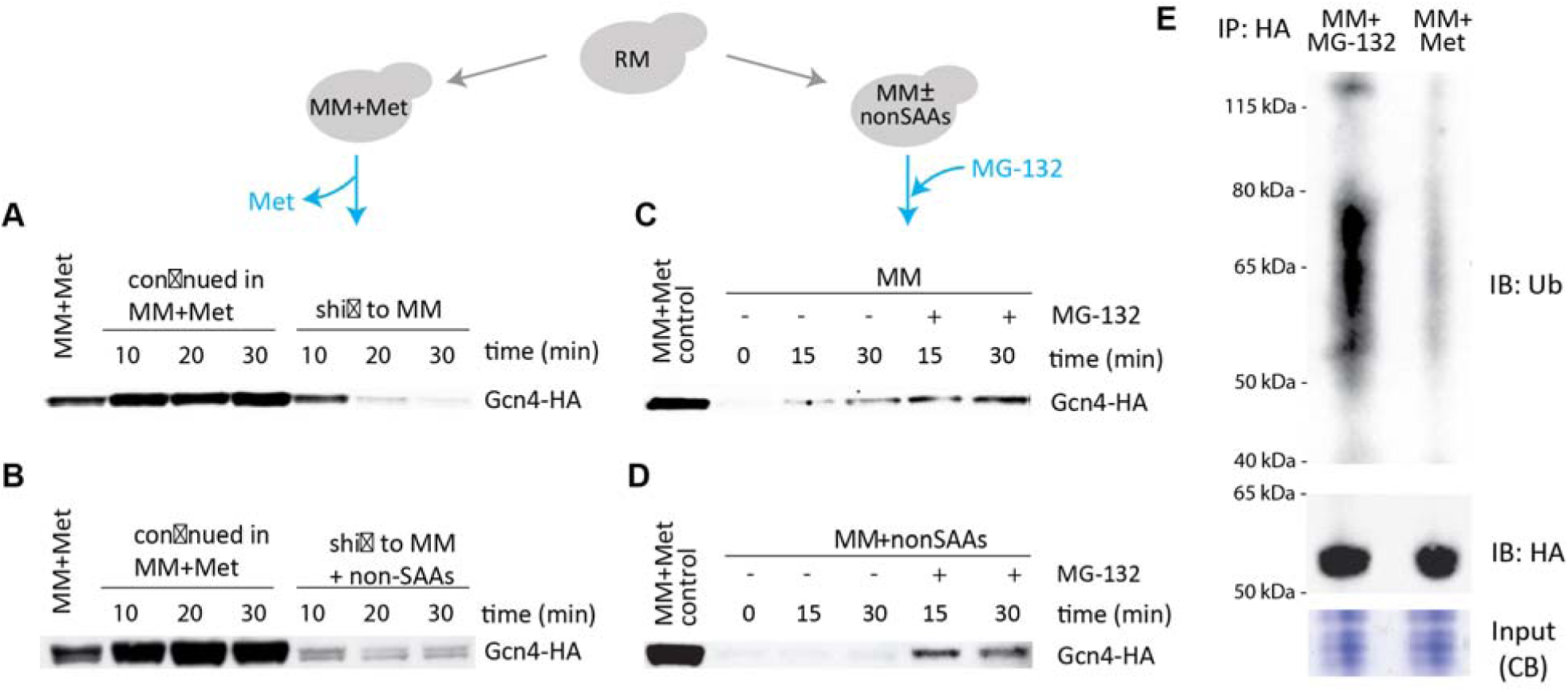
Methionine inhibits Gcn4 degradation. A) Cells growing in MM+Met were shifted to MM, and Gcn4 protein amounts were estimated across a short time course. Notably, Gcn4 amounts decrease substantially within 10 minutes of shift to MM. B) Cells growing in MM+Met were shifted to MM+ all non-sulfur amino acids (non-SAA), and Gcn4 protein amounts were estimated across a short time course. Notably, Gcn4 amounts decrease substantially within 10 minutes of shift to MM. C) Proteasomal degradation of Gcn4 in the absence of methionine. Cells growing in MM+Met were shifted to MM, in the presence or absence of the proteasomal inhibitor MG132, and Gcn4 protein amounts were estimated across a short time course. Gcn4 amounts clearly accumulate in the presence of MG132. D) Cells growing in MM+Met were shifted to MM+non-SAA, in the presence or absence of the proteasomal inhibitor MG132, and Gcn4 protein amounts were estimated across a short time course. Gcn4 amounts accumulate in the presence of MG132. E) Gcn4 ubiquitination decreases in the presence of methionine. Cells growing in MM (with MG132 added) or MM+Met were collected, and Gcn4 was immunopurified. The immunopurified Gcn4 was resolved on SDS-PAGE gels, and poly-ubiquitin chains detected by western blotting using an ubiquitin-specific antibody. Cells in MM+Met show substantially decreased polyubiquitin bands. IB: immunoblot, IP: immunopurification, CB: coomassie blue stain.

Next, we examined the ubiquitination status of Gcn4 and its modulation by methionine. Because ubiquitinated species are rapidly turned over by the 26S proteasome, we used MG-132 to inhibit degradation and accumulate polyubiquitin tagged proteins. Using a setup similar to earlier experiments, we initially adapted cells to MM+Met to allow Gcn4 to accumulate and then shifted them to MM in the presence of either methionine or MG-132. Particularly, immunoblot analysis showed a smear of high molecular weight, ubiquitinated forms only in Gcn4 immuno purified from cells shifted to media lacking methionine (Figure 3E). However, such ubiquitinated species were almost undetectable when Gcn4 was immuno purified from cells that had remained in MM+Met medium (Figure 3E). This suggests that although Gcn4 is constantly synthesized in MM regardless of methionine availability, abundant methionine specifically inhibits the proteasomal degradation of Gcn4.

### Methionine does not alter the composition of the Gcn4 targeting ubiquitination machinery

Gcn4 ubiquitination is known to be carried out by the Skp1-Cullin-F box (SCF) family of ubiquitin E3 ligases (42, 43) (Figure 4A). Skp1-Cullin are evolutionarily conserved proteins that form a stable complex, while the F-box is a variable component that defines the target specificity (44). Cdc4 is a F-box protein required for G1/S and G2/M cell cycle transitions. In addition to its role in the cell cycle, this protein ubiquitinates Gcn4 when part of the SCF^Cdc4^ complex (42, 43). Hence, we first examined whether methionine alters Cdc4 amounts, which could modulate ubiquitination activity towards substrates including Gcn4. We compared Cdc4 levels in cells grown in RM to that in cells after shift to MM or MM+Met. Cdc4 amounts did not change across these conditions tested (Figure 4B), suggesting that it is not specifically methionine regulated. We next asked if methionine instead alters the binding of Cdc4 to Skp1-Cullin (Cdc53), which will impact the functional SCF^Cdc4^ pool. We engineered a strain where both Cdc4 and Cdc53 were tagged with HA and FLAG epitopes at their respective chromosomal loci, grew these strains in RM and shifted these cells to MM and MM+Met before collection. We grew cells for 2h post-shift to allow sufficient time for the re-equilibration of cellular SCF complexes. Cdc53 was immunoprecipitated and the samples subject to HA immunoblots to assess the association between Cdc4 and Skp1-Cdc53 complex. Regardless of the presence of methionine, comparable levels of Cdc4 was immunoprecipitated in all conditions (Figure 4C). We obtained similar results when Cdc4 was immunoprecipitated instead, or if cells were shifted for only 20min using the setup described in Figure 3. These results suggest that methionine does not regulate the amounts of the ubiquitination complex known to ubiquitinate Gcn4.

**Figure 4:**
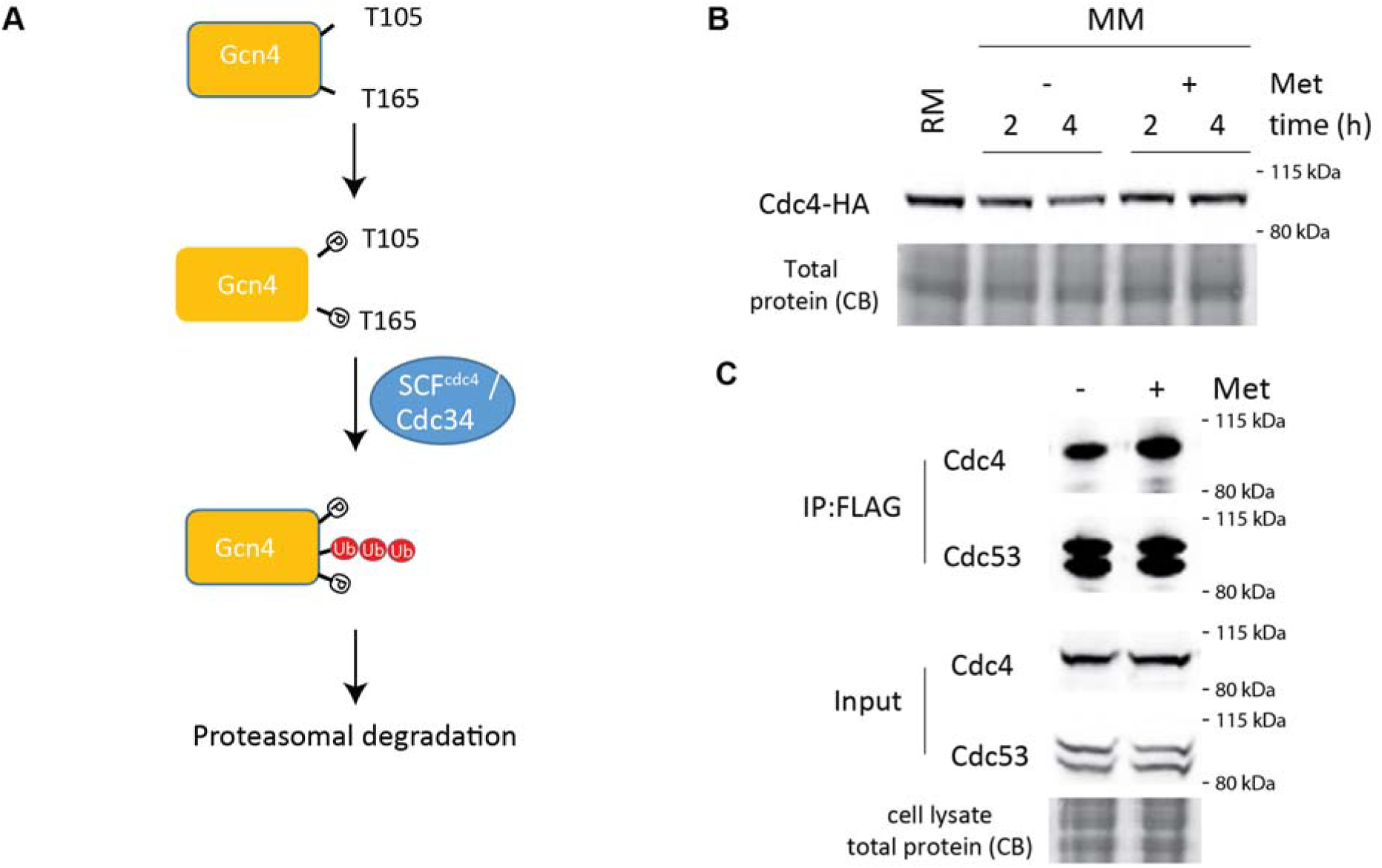
Methionine does not alter the composition of the Gcn4 targeting ubiquitination machinery. A) A schematic illustration indicating known mechanisms of Gcn4 degradation. Gcn4 can be phosphorylated (dependent on the Pho85 kinase), and this phosphorylation of two threonine residues leads to the ubiquitination of Gcn4 by the SCF (Cdc34) complex (with the specific F-box being Cdc4). This leads to Gcn4 degradation. B) Amounts of Cdc4 in MM or MM+Met. Cells in RM were shifted to MM or MM+Met and Cdc4 amounts were measured by Western blotting. C) Interaction of Cdc34, Cdc53 and Cdc4 and formation of the SCF complex. Cells in MM or MM+Met were collected, Cdc53 (FLAG) was immunoprecipitated and the samples subject to HA immunoblots to assess the association between Cdc4 and Skp1-Cdc53 complex.

### Phosphorylation at Thr165 is critical for Gcn4 degradation in the absence of methionine

The targeting of Gcn4 for ubiquitination by the SCF^Cdc4^ is regulated upstream by phosphorylation. In amino acid replete conditions, the cyclin dependent kinase (CDK) Pho85 in coordination with the cyclin Pcl5 phosphorylates Gcn4 (42, 45) (Figure 4A), and phosphorylated Gcn4 becomes a substrate of the SCF^CDC4^ ubiquitination complex. Therefore, we asked whether methionine modulates Pho85 or Pcl5 protein abundance. We compared Pho85 and Pcl5 protein in methionine replete or deplete medium, and found that these two proteins are present at similar amounts (Supplementary Figure 3), suggesting that kinase availability was not a limiting factor in putative Gcn4 phosphorylation in either condition. We then tested the importance of phosphorylation for Gcn4 stability in the absence or presence of methionine. We generated *pho85Δ* and *pcl5Δ* cells, and asked how the deletion of these proteins alters Gcn4 protein amounts. Strikingly, we found that the absence of either Pho85 or Pcl5 was sufficient to stabilize Gcn4 even in the absence of methionine (Figure 5A). We next directly investigated the kinetics of Gcn4 degradation upon removal of methionine, by shifting cells from RM to MM and measuring Gcn4 over a very brief time course. Here, we observed a rapid reduction in Gcn4 amounts in wildtype cells (Figure 5B), consistent with the kinetics observed in an earlier, different setup (Figure 3A). Contrastingly, in *pcl5Δ* cells, Gcn4 degradation was delayed, resulting in elevated protein levels even after 30 min of shift in MM without methionine. Further, the amount of Gcn4 in both *pho85Δ* and *pcl5Δ* cells in MM+Met was comparable and high. These data collectively indicated that Pcl5 kinase activity towards Gcn4 remains high in both methionine-replete and deplete medium. However, this also suggested that the loss of phosphorylation was sufficient to stabilize Gcn4 in the absence of supplemented methionine, indicating that Gcn4 degradation proceeds via the phosphorylation-ubiquitination pathway and this pathway is likely modulated by methionine.

**Figure 5:**
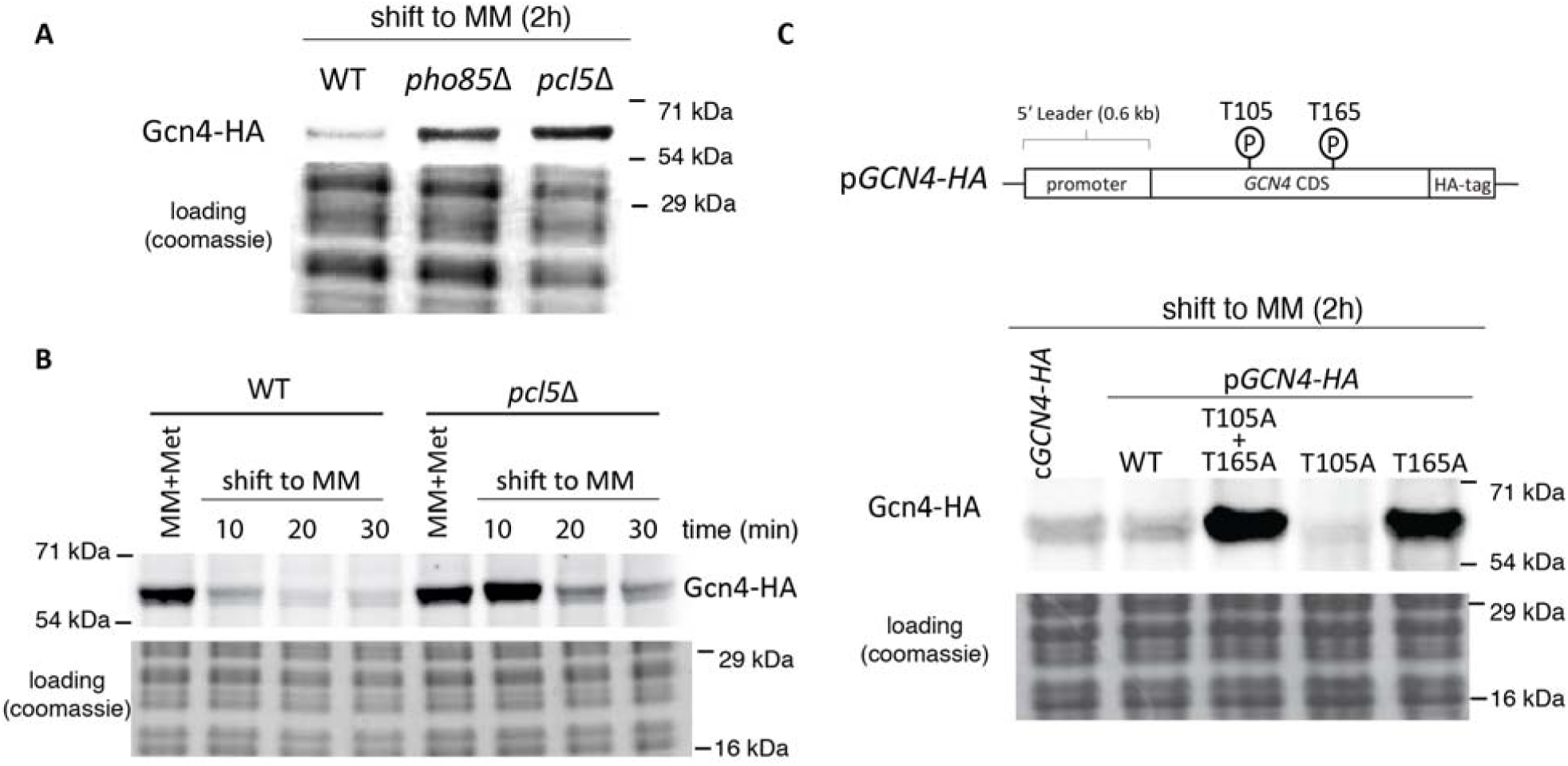
Phosphorylation at Thr165 is critical for Gcn4 degradation in the absence of methionine. A) Loss of phosphorylation stabilizes Gcn4 in MM. Wild-type, *pcl5Δ* and *pho85Δ* cells growing in RM were shifted to MM, and amounts of Gcn4 in these cells were estimated by Western blot. B) Loss of Gcn4 phosphorylation maintains Gcn4 amounts when cells are shifted to MM. Wild-type or *pcl5Δ* cells in MM+Met were shifted to MM, and the amounts of Gcn4 were estimated by Western blotting. C) Gcn4 phospho-mutants are stabilized in MM. The known phosphorylation residues (T105 and T165) of Gcn4 were mutated to alanines, and the amounts of Gcn4 in wild-type cells or T105A/T165A mutant Gcn4 cells were compared.

The Pho85/Pcl5 complex phosphorylates Gcn4 specifically at two independent sites, Thr105 and Thr165 (42, 43). To unambiguously study the phosphorylation dependent regulation of Gcn4 by methionine, we expressed native Gcn4 (with its endogenous promoter and regulatory regions) with a C-terminal HA tag, along with alanine-mutations of the respective threonine residues (T105 and T165) of Gcn4 individually or in combination, using a centromeric (CEN.ARS) plasmid (see Supplementary Figure 4). We expressed these forms of Gcn4 in wildtype yeast and examined Gcn4 levels after RM to MM shift. Wildtype Gcn4 expressed from the plasmid was degraded in an indistinguishable manner to chromosomally tagged protein, confirming that our system fully recapitulates our observations with native Gcn4 (Figure 5C). Notably, the double alanine mutant of Gcn4 was stabilized in MM (where methionine was not supplemented). Comparing the effects of the single mutants, the Thr105 single mutant was degraded similar to wildtype whereas the Thr165 mutant was stabilized (Figure 5C). These data conclusively show that phosphorylation of Gcn4 at Thr165 and subsequent ubiquitin-mediated degradation must be inhibited by methionine.

### The PP2A methyl transferase Ppm1 is required for methionine-mediated Gcn4 stabilization and function

We are therefore left with the observation that the ubiquitination and degradation of Gcn4 are high in the absence of methionine, Gcn4 is stabilized by methionine, the degradation of Gcn4 depends on known phosphorylation sites on this protein, but the known kinases itself appear to be unaffected by methionine. Thus, we wondered whether this stabilization due to methionine might instead be mediated by the selective activity of a protein phosphatase, which decreases the phosphorylation state of Gcn4. If, in the presence of methionine, a phosphatase specifically targets Gcn4 to dephosphorylate it, this would reduce Gcn4 phosphorylation (even in the presence of the kinase), and result in Gcn4 being stabilized. Interestingly, extensive evidence suggests important roles for a specific, methionine-responsive phosphatase in these conditions. Methionine strongly boosts the production of SAM, an important methyl donor (46), as established in these experimental conditions used (13, 47). Notably, these studies have shown that methionine/SAM drives the methylation of the catalytic subunit of the protein phosphatase, PP2A (13, 17). The methyltransferase, Ppm1p (called LCMT1 in mammals) specifically methylates PP2A at its carboxy-terminal leucine residue (48, 49). Further, methylated PP2A (via Ppm1 activity) preferentially dephosphorylates specific substrates such as the TORC1 repressor, Npr2, in a methionine-responsive manner (47). This tunable substrate preference of PP2A made us ask whether methylated PP2A and Ppm1 activity might regulate Gcn4 stability (Figure 6A). If this hypothesis is correct, an explicit prediction is that if PP2A methylation is prevented in the presence of methionine, Gcn4 will be destabilized and degraded.

**Figure 6:**
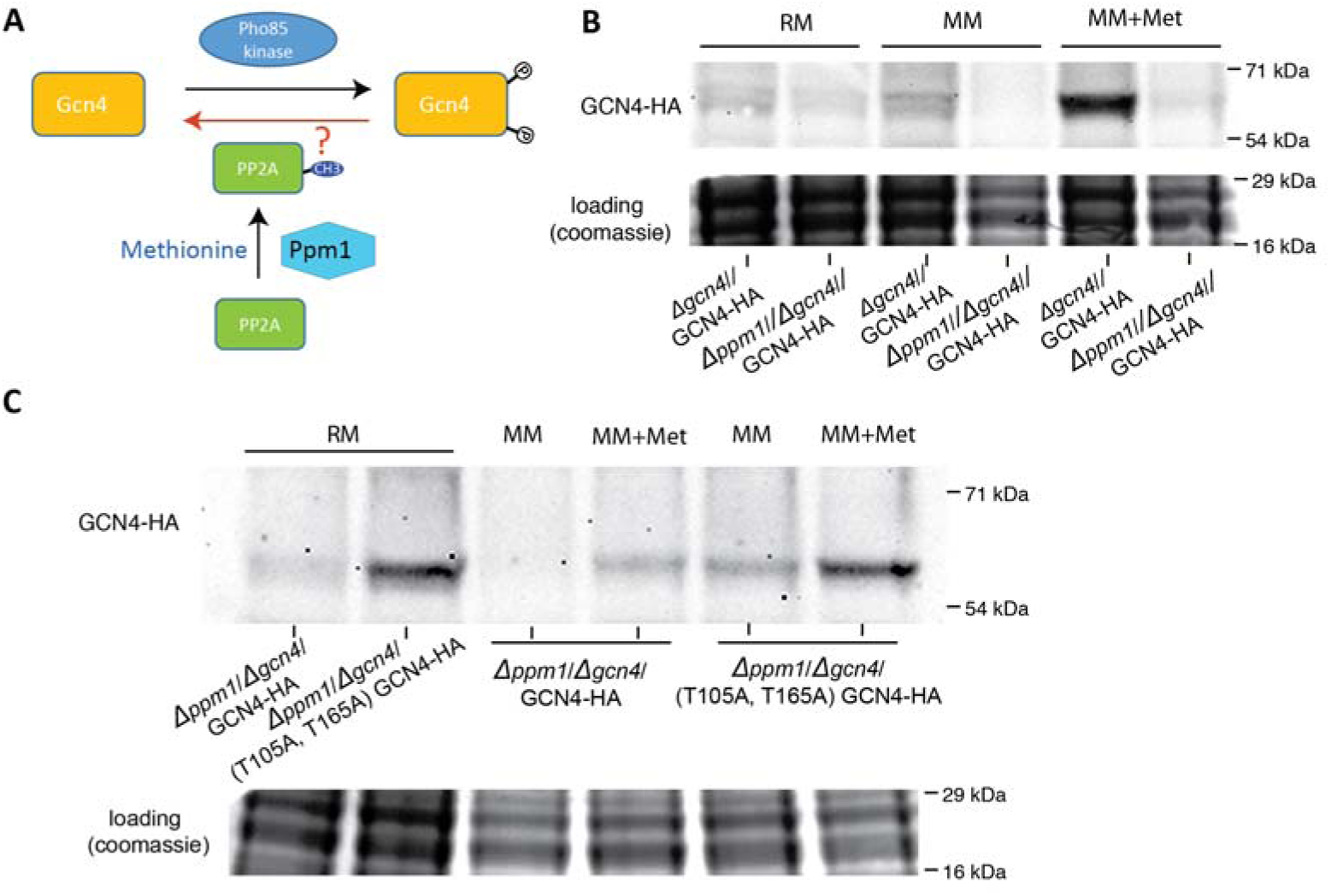
The PP2A methyl transferase Ppm1 is required for methionine-dependent Gcn4 stabilization and function. A) A hypothetical mode by which Gcn4 phosphorylation might be regulated. In methionine sufficiency, PP2A is carboxyl-methylated by the methyltransferase Ppm1. If methylated PP2A targets Gcn4 for dephosphorylation, this will decrease Gcn4 ubiquitination and therefore degradation. B) Loss of Ppm1 decreases Gcn4 amounts in MM+Met. The amounts of Gcn4 protein were compared in wild-type and *ppm1Δ* cells in both MM and MM+Met by Western blotting. A representative gel from at least three experiments is shown. Also see Supplemental Figure 5 for Gcn4 amounts in PP2A(L->A) cells. C) Gcn4 phospho-mutants are stabilized in *ppm1Δ* cells in MM+Met. Gcn4 or Gcn4(T105A/T165A) protein amounts were compared in *ppm1Δ* cells in MM or MM+Met, as indicated.

We directly tested this surmise using a strain lacking the Ppm1 methyltransferase. We examined Gcn4 levels after a shift to MM or MM+Met, in wildtype and *ppm1Δ* cells. After shifting cells to methionine-deplete medium (MM), Gcn4 amounts substantially decrease in both strains, indicating that the Gcn4 degradation pathway remained functional in the mutant. Strikingly, in the *ppm1Δ* strain, we observed substantially reduced amounts of Gcn4 even in the presence of methionine, compared to wild-type cells (Figure 6B). This indicates that Ppm1 activity is necessary to maintain high Gcn4 protein, in the presence of methionine. We next directly tested the importance of Gcn4 phosphorylation in this Ppm1 dependent degradation. We introduced the phosphorylation-insensitive point mutants of Gcn4, and asked if this restored the Gcn4 levels when methionine is present, and Ppm1 is absent. We expressed either wildtype Gcn4 or the Thr105Ala/Thr165Ala stabilization mutants in wildtype and *ppm1Δ* cells, and observed that in *ppm1Δ* cells, the amounts of the alanine-mutant form of Gcn4 remained high upon shift to MM+Met medium, whereas wild type Gcn4 protein showed a striking reduction in amounts (Figure 6C). These data reveal that Ppm1 activity, which is required for PP2A carboxy-methylation, is critical for the stabilization of Gcn4. We further reiterated these results using a yeast strain where the two isoforms of the PP2A catalytic subunit (Pph21 and Pph22) both have their respective C-terminal leucine residues mutated to alanines (and hence can no longer be methylated) (13). Consistent with the results observed with the *ppm1Δ* strain, in a PP2A methylation-deficient strain (PP2A-L->A), Gcn4 protein was not detectable in cells shifted to MM+Met (Supplementary Figure 4). Taken together, these observations firmly establish that reversal of Gcn4 phosphorylation by methionine sensitive Ppm1 and therefore PP2A phosphatase activity is necessary and sufficient to rescue Gcn4 from ubiquitin mediated degradation.

### Ppm1/PP2A-methylation is critical for the Gcn4 mediated anabolic response in methionine replete conditions

Finally, we addressed the functional consequences of loss of PP2A methylation on Gcn4 mediated outputs, when methionine is abundant. We compared relative transcript amounts of select direct Gcn4-targets (22) by qRT-PCR analysis, in WT, Pph21/22L→A, *ppm1Δ* and *gcn4Δ* (control) cells. Expectedly, in the Pph21/22L→A and *ppm1Δ* cells, Gcn4 target transcripts were significantly reduced compared to WT cells (Supplementary Figure 5). We next directly analyzed the metabolic consequences in Pph21/22L→A and *ppm1Δ* cells, in the presence of methionine. For this we estimated the *de novo* synthesis of key Gcn4 dependent metabolites by estimating the incorporation of stable isotopes of nitrogen into nucleotides or amino acids after pulse-labeling, using LC-MS/MS (liquid chromatography-mass spectrometric) analysis (see schematic of Figure 7A) (see materials and methods for the monitored mass transitions). The results (Figure 7A) unambiguously revealed that Pph21/22L→A, *ppm1Δ* and *gcn4Δ* cells all show comparable, poor label incorporation into these newly synthesized metabolites when compared to the WT cells.

**Figure 7:**
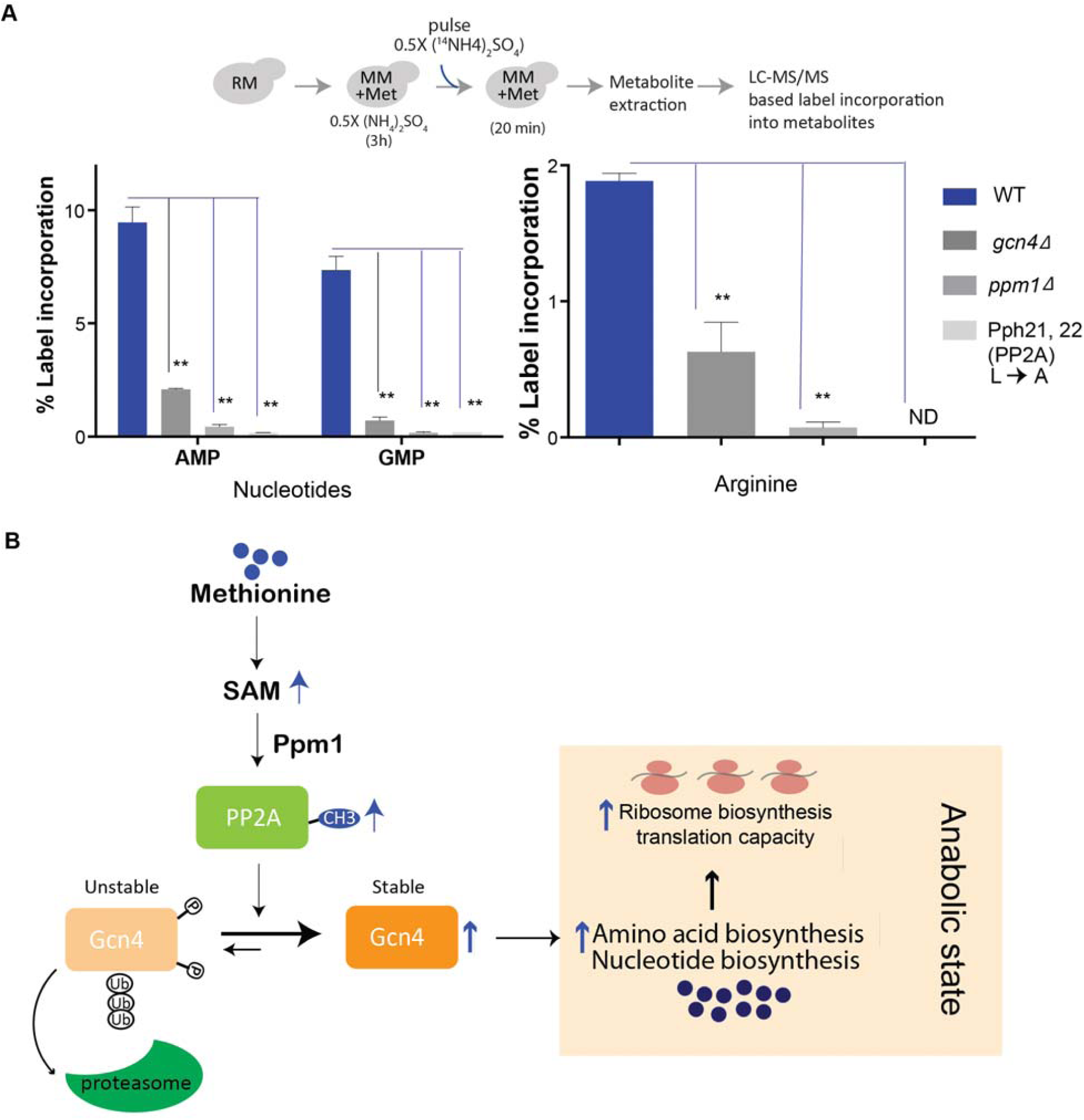
Ppm1/PP2A-methylation is critical for the Gcn4 mediated anabolic response in methionine replete conditions. A) Gcn4 dependent anabolic precursor synthesis is absent in *ppm1Δ* and PP2A(L->A) cells. Quantitative, targeted LC-MS/MS based analysis of the *de novo* synthesis of Gcn4-dependent metabolites (AMP, GMP and arginine). The indicated cells were shifted to MM+Met, pulsed with ^14^N-labelled ammonium sulfate, metabolites extracted, and label-incorporation into the respective newly synthesized metabolites were estimated. B) Model illustrating how Gcn4 is stabilized by methionine, via the action of Ppm1/PP2A-methylation. In sufficient methionine, S-adenosyl methionine amounts are high, and therefore the methyltransferase Ppm1 methylates the carboxyl-terminal leucine of PP2A. Methylated PP2A preferentially dephosphorylates Gcn4, thereby altering Gcn4 towards a less phosphorylated state. This decreased phosphorylation prevents the ubiquitination and therefore degradation of Gcn4, resulting in its accumulation. High Gcn4 amounts now drive amino acid and nucleotide biosynthesis, enabling cells to maintain an anabolic program.

Taken together, these data show that the Ppm1 mediated methylation of PP2A controls the stability of Gcn4, and thereby the methionine-mediated anabolic program controlled by Gcn4.

## Discussion

In this study we elucidate a regulatory mechanism by which methionine induces Gcn4, in order to drive a growth program. In methionine replete conditions, cells stabilize Gcn4, via the Ppm1-dependent methylation of the major protein phosphatase PP2A. The methylated form of PP2A preferentially shifts the pool of Gcn4 towards a dephosphorylated state. Dephosphorylated Gcn4 is protected from ubiquitination and subsequent proteasomal degradation, and thereby increases in amounts. Ppm1 activity and PP2A carboxymethylation dependent stabilization of Gcn4 is essential for the sustained increase in amino acid and nucleotide synthesis in the presence of methionine. Our findings therefore reveal a fundamental, metabolic function of PP2A, which is to increase the availability of key anabolic precursors (amino acids and nucleotides), upon sufficient availability of methionine (via its metabolite s-adenosylmethionine). This is achieved by directly regulating the amounts of a metabolic master-regulatory transcription factor, Gcn4. This mechanistic model is illustrated in Figure 7B.

It becomes apparent that the mechanisms by which Gcn4 protein is regulated depends on context and metabolic state of the cell. The best understood mechanism of Gcn4 (or ATF4) activation is via increased Gcn4 translation during amino acid starvation (24, 50). In conditions of acute amino acid starvation (including methionine starvation), cells activate the Gcn2 kinase and eIF2α phosphorylation, which reduces overall translation while increasing Gcn4 translation due to the unique regulatory elements in Gcn4 mRNA. Through this mechanism, cells restore amino acid amounts during starvation. In contrast, here we illustrate an alternate mode of increasing Gcn4 activity when cells are in a *growth state*. Here, Gcn4 accumulates due to its increased stability in the presence of abundant methionine. While the degradation of Gcn4 due to phosphorylation dependent ubiquitination is known (42, 43), the physiological contexts where this mechanism is important are poorly studied. Here, we show how, by using distinct signaling modules to increase Gcn4 stability, cells might activate this protein in a context-dependent manner. For example, in an amino acid replete context, if cells need to degrade Gcn4, it is easy to imagine the activation of specific kinases that phosphorylate Gcn4, leading to its degradation. However, alternate scenarios when cells are in a growth state but still require Gcn4 function become easily possible. Methionine is a growth signal, activating both translation and the biosynthesis of metabolic precursors required to sustain growth (11, 14). This synthesis of metabolic precursors is Gcn4 dependent, and by using this distinct, methionine-responsive phosphatase based mechanism, cells can increase Gcn4 amounts despite high rates of protein translation. By reducing the phosphorylation of Gcn4 in a methionine-dependent manner, Gcn4 can be selectively stabilized, to drive anabolism.

A central concept emerging from this study is how metabolite responsive signaling can directly control metabolism. How methionine regulates signaling processes that directly control metabolic outputs is not yet clear. While methionine (via SAM) activates the TORC1 (13, 15, 16), this mechanism does not explain how Gcn4 is activated, and how methionine controls the metabolic program. However, the regulation of Gcn4 phosphorylation via a methionine/SAM specific modification of a phosphatase now provides a mechanism of action. The evidence for regulated PP2A activity and substrate selectivity via carboxyl-methylation is now extensive (48, 49, 51–56). The carboxyl-methylation of PP2A, by Ppm1/LCMT1, alters the substrate binding preference or specific activity of this phosphatase, as established in numerous systems (48, 51, 54, 55). What has been poorly explored are the contexts in which this methylation-dependent regulation plays a role. We now establish that the methylated form of PP2A (which depends on the methyltransferase Ppm1) directly regulates cellular metabolic state in response to methionine/SAM, by increasing Gcn4 stability. This study therefore completes an important conceptual loop: starting from the metabolite (methionine), a specific signaling output that depends on the metabolite is regulated (the SAM dependent methylation of PP2A), which increases the dephosphorylation of Gcn4, and this in turn controls the overall metabolic state of the cell. This is an effective, directed means by which the cell can prioritize and manage available resources to commit to a growth state.

As introduced earlier, Gcn4 is an evolutionarily conserved regulator of amino acid biosynthesis. Although it is primarily studied in the context of starvation, during a methionine driven growth program Gcn4 activity fuels the synthesis of required biosynthetic precursors to drive growth. Interestingly, such scenarios are also observed in several tumors, where the mammalian ortholog of Gcn4 (ATF4) is induced and helps in tumor proliferation (57–60). The mechanisms of induction of ATF4 in these contexts have not been well studied, and the mechanism observed in this study is plausible in those contexts of high growth.

We summarize a unifying model for how methionine launches a multi-pronged transcriptional and metabolic program to drive growth (11). The key elements in this transformation include tRNA thiolation mediated routing of carbon-flux towards the PPP as well as maintaining overall metabolic homeostasis (41), Gcn4-mediated increased biosynthesis of amino acids and nucleotides (14, 22), maintaining translation capacity (22), balancing phospholipid and histone methylation (17), and SAM-mediated increased translation via the activation of the TORC1 pathway (13, 15, 16). The role of the methyltransferase Ppm1, and methylation dependent PP2A activity in the methionine-induced growth program appears to be central, since methylated PP2A activates the TORC1 pathway (13), global histone methylation (17, 61), and controls anabolic precursor synthesis via the activation of Gcn4 (as seen in this study). Given the evolutionary conservation of all these processes observed in yeast and mammals, including the degradation of ATF4 via a phosphorylation dependent interaction with a ubiquitin ligase (62), it will be interesting to evaluate the roles of methylated PP2A dependent signaling outputs in relevant contexts of growth regulation in healthy and diseased tissue. SAM-sensitive methylation of PP2A could be a universal mode by which cells directly sense this key metabolite (methionine), dictate global phosphorylation status and correspondingly regulate metabolism to drive growth programs.

## Materials and methods

### Yeast strains and growth media

The prototrophic CEN.PK strain (referred to as wild type, WT) was used in all experiments (*van Dijken et al., 2000*). Strains with gene deletions or chromosomally tagged proteins (at the C-terminus) were generated as described elsewhere (ref). Strains used in this study are listed in **Table 1**.

**Table 1:** Strains used in this study.

| Strain background | Genotype | Reference |
| --- | --- | --- |
| CEN.PK | MAT a (full prototroph) | From (63) |
| CEN.PK | MAT a *GCN4-HA-KanMX* | From (14) |
| CEN.PK | MAT a *gcn4Δ::NAT* | From (14) |
| CEN.PK | MAT a *SUI2-FLAG-NAT* | This study |
| CEN.PK | MAT a *SUI2-FLAG-NAT GCN4-HA-KanMX* | This study |
| CEN.PK | MAT a *SUI2-FLAG-NAT GCN4-HA-KanMX gcn2Δ::HYG* | This study |
| CEN.PK | MAT a *GCN4-HA-KanMX pdr5Δ::HYG* | This study |
| CEN.PK | MAT a *CDC4-HA::KanMX* | This study |
| CEN.PK | MAT a *CDC4-HA::KanMX CDC53-FLAG::Hyg* | This study |
| CEN.PK | MAT a *pho85Δ::NAT gcn4Δ::KanMX* | This study |
| CEN.PK | MAT a *pcl5Δ::NAT gcn4Δ::KanMX* | This study |
| CEN.PK | MAT a *ppm1Δ::NAT gcn4Δ::Hyg* | This study |
| CEN.PK | MAT a *ppm1Δ::NAT GCN4-HA::KanMX* | This study |
| CEN.PK | MAT a *pdr5Δ::Hyg ppm1Δ::NAT GCN4-HA::KanMX* | This study |
| CEN.PK | Pph21/22L→A | From (13) |

The growth media used in this study are RM (1% yeast extract, 2% peptone and 2% lactate) and MM (0.17% yeast nitrogen base without amino acids, 0.5% ammonium sulfate and 2% lactate). All amino acids were supplemented at 2 mM. NonSAAs refers to the mixture of all standard amino acids (2 mM each) except methionine, cysteine and tyrosine.

The indicated strains were grown in RM with repeated dilutions (∼36 hours), and the culture in the log phase (absorbance at 600 nm of ∼1.2) was subsequently switched to MM, with or without addition of the indicated amino acids.

### Polysome analysis

The cells (50 OD) were treated with cycloheximide (100 μg/ml) for 15 minutes before harvesting in ice-filled, cold centrifuge bottles. The pellet obtained after centrifugation (8000g, 5 min, 4°C) was once washed, and then re-suspended in 0.5 ml lysis buffer (10 mM Tris-HCl pH 7, 0.1 M NaCl, 30 mM MgCl_2_, 1U/μl Ribolock RNase inhibitor and 80 μg/ml cycloheximide) with chilled, baked acid-washed glass beads. After bead-beating, 0.5 ml lysis buffer was added and the crude extract obtained was centrifuged (18407g, 10 min, 4°C) and the supernatant (200 μl) containing polysomes is fractionated on a sucrose gradient (7%-47% sucrose; containing 20 mM Tris-HCl pH 7, 140 mM KCl, 5 mM MgCl_2_, 1 mM DTT and 50 μg/ml cycloheximide) by ultra-centrifugation (35000 rpm, 3 h, 4°C). The polysome profile was obtained on an ISCO fractionator, wherein the absorbance at 254 nm was traced. For polysome: monosome ratio calculation, area under the curve was quantified using ImageJ software (*Rasband, ImageJ*, U. S. National Institutes of Health, Bethesda, Maryland, USA, http://imagej.nih.gov/ij/, 1997-2016).

### Western blot

Approximately ten OD cells were collected from respective cultures, pelleted and flash-frozen in liquid nitrogen until further use. The cells were re-suspended in 400 μl of 10% trichloroacetic acid and lysed by bead-beating three times: 30 sec of beating and then 1 min of cooling on ice. The precipitates were collected by centrifugation, re-suspended in 400 μl of SDS-glycerol buffer (7.3% SDS, 29.1% glycerol and 83.3 mM Tris base) and heated at 100°C for 10 min. The supernatant after centrifugation was treated as the crude extract. Protein concentrations from extracts were estimated using bicinchoninic acid assay (Thermo Scientific). Equal amounts of samples were resolved on 4 to 12% Bis-Tris gels (Invitrogen). Coomassie blue–stained gels were used as loading controls. Western blots were developed using the antibodies against the respective tags. We used the following primary antibodies: monoclonal FLAG M2 (Sigma), HA (12CA5, Roche), and phosphor-eIF2α Ser51 (9721, Cell Signaling). Horseradish peroxidase– conjugated secondary antibodies (mouse and rabbit) were obtained from Sigma. For Western blotting, standard enhanced chemiluminescence reagents (GE Healthcare) were used. ImageJ was used for quantification.

### *GCN4-Luc* constructs

A series of constructs having variations of the *GCN4* promoter were generated as shown in Supplementary Figure 1 and Figure 2B. The luciferase reporter gene synthesized into a pGL3basic vector (Promega), and sub-cloned in *EcoR*I and *Xho*I sites of the CEN/ARS, single-copy plasmid p417-TEF-KAN plasmid (modified from p417-CYC, MoBiTech, Germany). The resulting plasmid was utilized for inserting different synthesized fragments (GeneArt), having indicated point mutations in the promoter elements and the first 55 codons of *GCN4* ORF, using *Sac*I and *EcoR*I sites just upstream of the luciferase gene. Thus, in all the constructs, *GCN4* promoter was followed by the *GCN4* ORF (encoding the first 55 amino acids) fused to the firefly luciferase coding sequence. The luciferase activity was measured using a luciferase assay kit (Promega, E1500).

### Gcn4 phospho-mutants

The complete uORF reading frame of *GCN4*, along with the Gcn4 coding sequence, followed by 6x-HA epitope tag at the carboxy terminus, followed by an Adh1 terminator sequence were amplified from genomic DNA obtained from the Gcn4-HA strain. This was cloned into a CEN.ARS plasmid (p417-cyc), where the plasmid promoter sequence was also replaced with the complete *GCN4* upstream regulatory region. This resulted in a plasmid that can express full length Gcn4 with a C-terminal 6HA epitope tag, with fully endogenous promoter and ORF regulatory regions included. This construct sequence was confirmed by sequencing. The identified phosphorylation site residues (T105 and T165) were mutated to alanine residues individually or in combination, by standard site-directed mutagenesis.

### RNA isolation and RT-qPCR

Total RNA from yeast cells was extracted using hot acid phenol method (*Collart and Oliviero, 2001*). SuperScript II reverse transcriptase (Invitrogen) was used for reverse transcription of the total RNA (1 μg). Quantitative PCR was performed with the synthesized cDNA using SYBR Green (Thermo scientific) and specific primers. *ACT1* was used for normalization of the transcript abundance. The primers used were *GCN4*: TGCTTACAACCGCAAACAGC and GCACGTTTTAGAGCAGCAGG, *ACT1*: TCGTTCCAATTTACGCTGGTT and CGGCCAAATCGATTCTCAA.

### Metabolite extractions and measurements by LC-MS/MS

For detecting ^15^N-label incorporation in amino acids and nucleotides, ^15^N-ammonium sulfate with all nitrogens labelled (Sigma-Aldrich) was used. At the end of the incubation with the spiked label, cells were rapidly harvested and metabolite extracted as described earlier (Wor 2018). Metabolites were measured using LC-MS/MS methods developed earlier (Walvekar et al, WOR, 2018). Standards were used for developing multiple reaction monitoring (MRM) methods on Sciex QTRAP 6500. All the measurements were done in the positive polarity mode. For all the nucleotide measurements, release of the nitrogen base was monitored. All the mass transitions of parent/product masses measured for the different metabolites are enlisted in **Table 2**.

**Table 2:** Mass transitions used for the ^15^N labeling experiment.

| Molecule of interest | Formula | Parent/Product (positive polarity) | Comment |
| --- | --- | --- | --- |
| Arg | C<sub>6</sub>H<sub>14</sub>N<sub>4</sub>O<sub>2</sub> | 175.2 / 60 | Product has only one N |
| <sup>15</sup>N_Arg_1 |  | 176.2 / 61 |  |
| <sup>15</sup>N_Arg_2 |  | 177.2 / 61 |  |
| <sup>15</sup>N_Arg_3 |  | 178.2 / 61 |  |
| <sup>15</sup>N_Arg_4 |  | 179.2 / 61 |  |
| AMP | C<sub>10</sub>H<sub>14</sub>N<sub>5</sub>O<sub>7</sub>P | 348/136 | Product has all N |
| <sup>15</sup>N_AMP_1 |  | 349/137 |  |
| <sup>15</sup>N_AMP_2 |  | 350/138 |  |
| <sup>15</sup>N_AMP_3 |  | 351/139 |  |
| <sup>15</sup>N_AMP_4 |  | 352/140 |  |
| <sup>15</sup>N_AMP_5 |  | 353/141 |  |
| GMP | C<sub>10</sub>H<sub>14</sub>N<sub>5</sub>O<sub>8</sub>P | 364/152 |  |
| <sup>15</sup>N_GMP_1 |  | 365/153 |  |
| <sup>15</sup>N_GMP_2 |  | 366/154 |  |
| <sup>15</sup>N_GMP_3 |  | 367/155 |  |
| <sup>15</sup>N_GMP_4 |  | 368/156 |  |
| <sup>15</sup>N_GMP_5 |  | 369/157 |  |

## Acknowledgements

We acknowledge extensive use of the NCBS/inStem/CCAMP mass spectrometry facilities for HPLC and mass spectrometry. AW, RG and RS acknowledge support from SERB National Postdoctoral Fellowships (PDF/2015/000225, PDF/2016/000416, and PDF/2016/001877), DST, Govt. of India. GK was supported by an inStem/NCBS Campus Fellowship and BT/IN/DBT-MRC (UK)/13/SM/2015-16. SL acknowledges support from a DBT-Wellcome Trust India Alliance Intermediate Fellowship (IA/I/14/2/501523), and intramural support.

## Notes

### Competing Interest Statement

The authors have declared no competing interest.

