## Supplementary Figures for "Methylated PP2A stabilizes Gcn4 to enable a methionine-induced anabolic program"

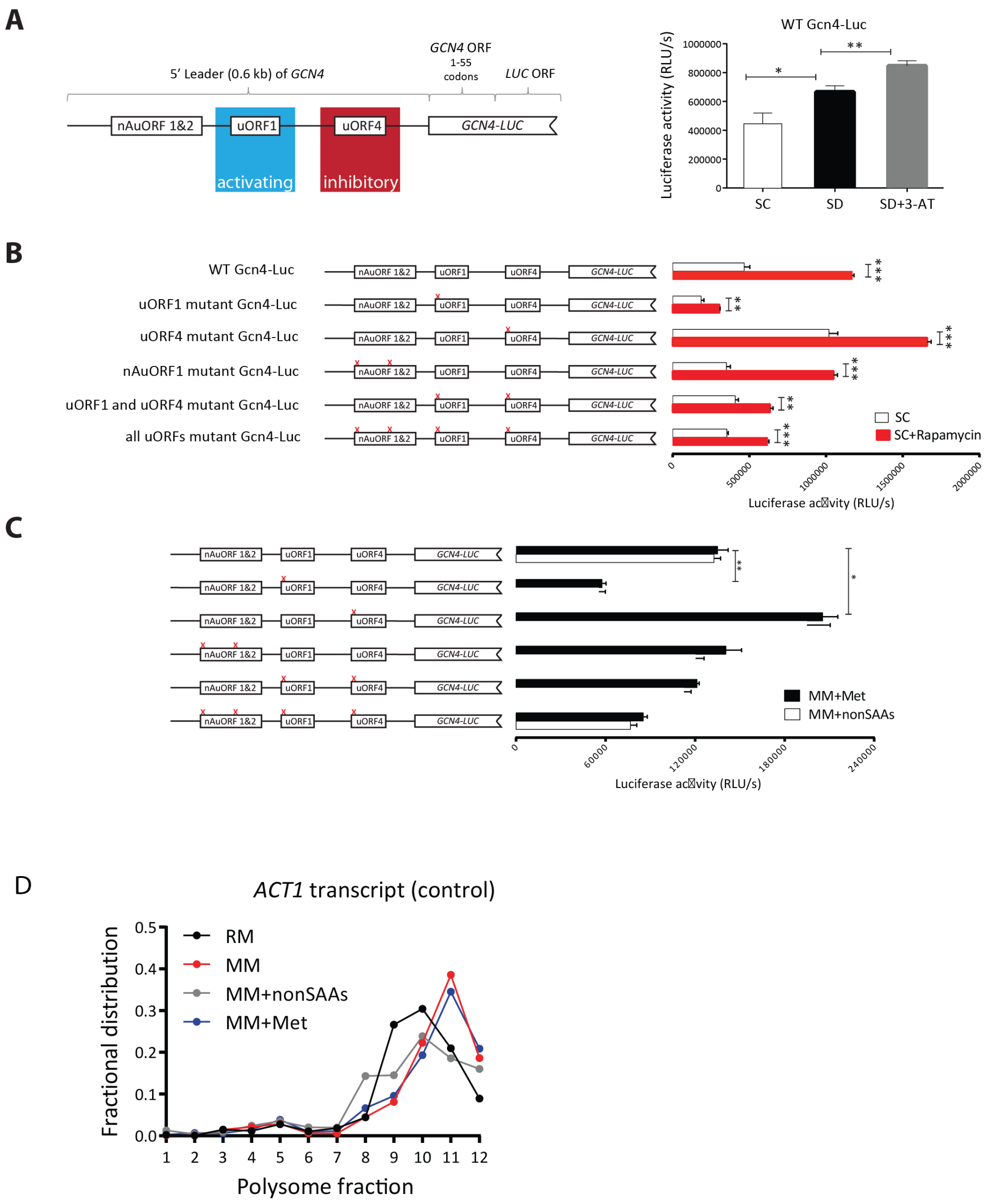


**Supplementary Figure 1: Gcn4 translation in MM, MM+nonSAAs and MM+Met, based on reporter activity or polysome fractions.**

A) Schematic illustrating the design of a luciferase based reporter to estimate the translation of *GCN4* transcripts, and validation of the reporter for *GCN4* translation in different medium.
B) Relative *GCN4* translation in complete minimal glucose medium with all amino acids (SC), or the same medium with rapamycin added to induce *GCN4* translation, as measured using a series of luciferase-based *GCN4* translation reporters. The relative luciferase activity is shown on the y-axis of the plots, while the different reporters used are illustrated on the left. The data shown are from three biological replicates, mean ± SD. * $p<0.05$ (Student's t-test).
C) Relative *GCN4* translation in MM+nonSAAs or MM+Met, as measured using a series of luciferase-based *GCN4* translation reporters. The relative luciferase

activity is shown on the y-axis of the plots, while the different reporters used are illustrated on the left. The data shown are from three biological replicates, mean ± SD. * $p<0.05$ (Student's t-test). MM+nonSAAs is MM supplemented with all non-sulfur containing amino acids (i.e. excluding methionine and cysteine).
D) *ACT1* transcript amounts in different polysome fractions, obtained from cells grown in RM, MM or MM+met. Transcripts were measured using standard, quantitative RT-qPCR approaches.

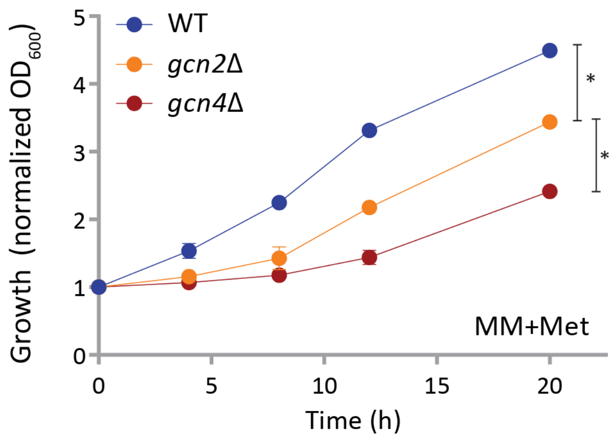


**Supplementary Figure 2: Comparison of growth in MM+Met**
Growth curves of wild type (WT), *gcn2*Δ and *gcn4*Δ cells, shifted from RM to MM+Met.

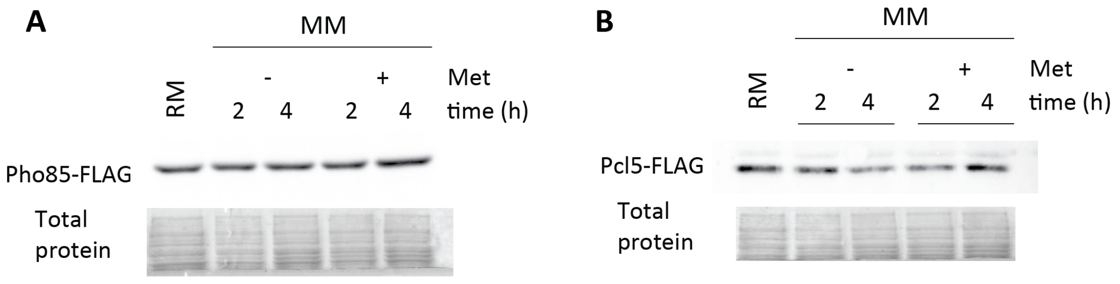


**Supplementary Figure 3: Comparison of relative amounts of the Pho85 and Pcl5 proteins, which comprise the Pho85-Pcl5 kinase comples.**
A) Amounts of Pho85 in MM or MM+Met. Cells in RM were shifted to MM or MM+Met and Pho85 amounts were measured by Western blotting (anti-FLAG).
B) Amounts of Pcl5 in MM or MM+Met. Cells in RM were shifted to MM or MM+Met and Pcl5 amounts were measured by Western blotting (anti-FLAG).

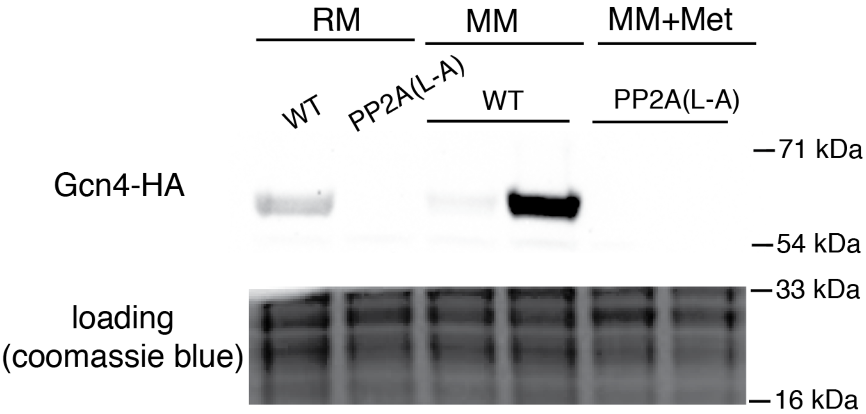


**Supplementary Figure 4: Gcn4 protein in WT and PP2A (Pph21/22) L→A cells.**

Loss of PP2A methylation decreases Gcn4 amounts in MM+Met. The amounts of Gcn4 protein were compared in wild-type and PP2A(L-A) cells in both MM and MM+Met by Western blotting. A representative gel from multiple experiments is shown.

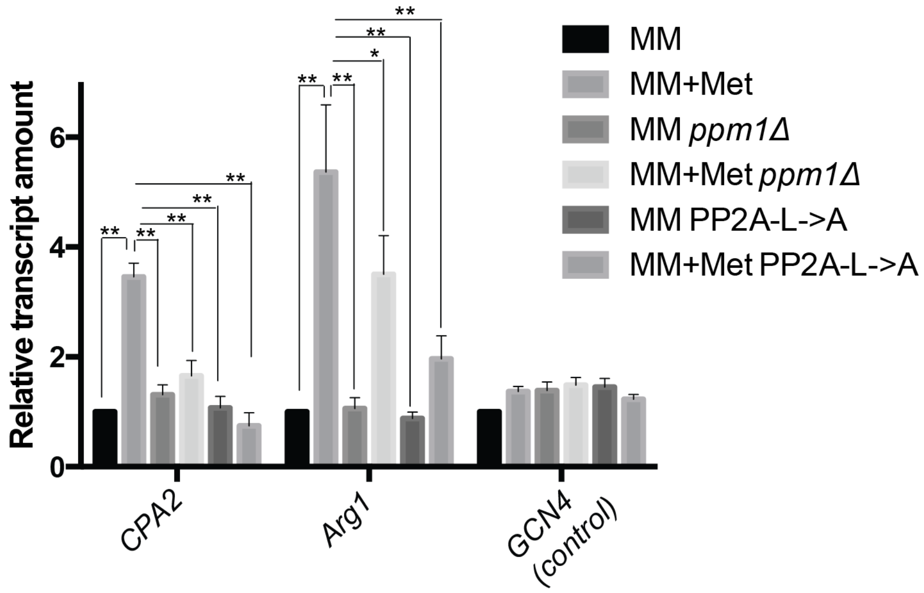


**Supplementary Figure 5: Relative transcript amounts of Gcn4 target genes in the indicated medium, and genetic background. *GCN4* transcripts are also indicated (as a control).**
